# Eliminating the type I restriction endonuclease from *Pseudomonas aeruginosa* PAO1 for optimised phage isolation

**DOI:** 10.1101/2025.05.30.656992

**Authors:** Ellie J. Tong, Kate A. Bickerton, Alina J. Creber, Steven L. Porter, Ben Temperton

## Abstract

Phage therapy is a promising treatment for multidrug-resistant bacterial infections. Due to their high host specificity, phages must be matched to the target clinical strains. Efficiently identifying appropriate phages and producing sufficient titres for clinical use requires comprehensive phage libraries and multiple propagation hosts. An idealised system would use a highly promiscuous bacterial host to isolate a broader range of phages and streamline optimised phage production. Anti-phage defences constrain bacterial host promiscuity, such as restriction-modification systems that recognise and cleave foreign DNA. Here, the type I restriction endonuclease, HsdR, was deleted from *Pseudomonas aeruginosa* PAO1 to make a more promiscuous phage isolation and propagation host. Removal of this endonuclease more than doubled the efficiency of phage propagation and yielded seven times more phages from freshwater samples than wildtype PAO1 – an important step in producing an optimised *P. aeruginosa* strain for isolating and propagating phages for clinical phage therapy.

**Graphical abstract:** (Created in BioRender.com) **[A]** The type I R-M system comprises three subunits HsdS, HsdM and HsdR. One HsdS and two HsdM subunits form the methyltransferase, and the addition of two HsdR subunits to this complex forms the restriction endonuclease. **[B]** In the wildtype PAO1, the type I R-M system destroys phage DNA while protecting the host. The HsdS subunit of the restriction endonuclease binds to unmethylated recognition sequences. Then, HsdR translocates the DNA, pulling it together in both directions until it collides with another restriction endonuclease, cleaving the DNA and preventing phage proliferation. The methyltransferase protects the host DNA through the methylation of recognition sequences, preventing the restriction endonuclease from binding. **[C]** In Δ*hsdR*, the restriction endonuclease cannot form, preventing phage DNA cleavage and increasing phage proliferation. The methyltransferase is still active, so both host and phage DNA are methylated, providing phage progeny with protection from R-M systems in future hosts.

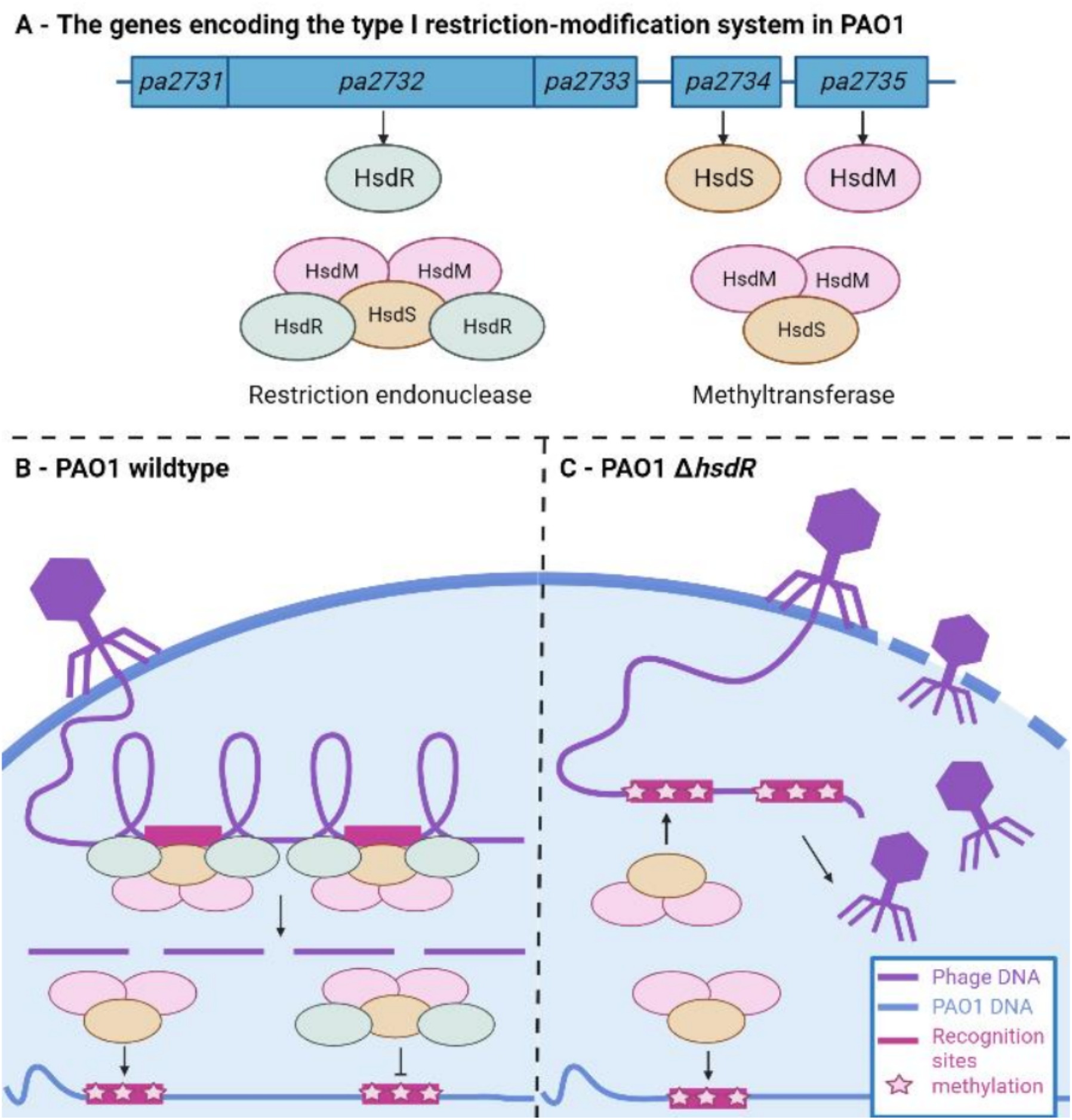

## Introduction

*Pseudomonas aeruginosa* is the fourth leading cause of hospital-acquired infections, with high morbidity and mortality in immunocompromised patients with burn wounds, sepsis, traumas, and respiratory conditions, including chronic obstructive pulmonary disease (COPD) and cystic fibrosis (CF)^1,2^. The World Health Organisation (WHO) considers *P. aeruginosa* a critical priority for new antibiotics, with 50% of intensive care isolates displaying multi-drug resistance^3,4^. The stalled development of novel antibiotics means alternative treatment strategies are required^5^.

Phage therapy represents a promising solution, where infections are treated with lytic bacteriophages (phages), the natural predators of bacteria^6^. Phages are highly specific, allowing targeted killing of the pathogen without damaging commensal microbes^7^.However, this specificity also requires phages to be matched to clinical isolates^8^. Comprehensive libraries of characterised phages enable efficient selection of specific phages for patients^9^. Building sufficiently broad phage libraries to match circulating pathogenic strains requires multiple and/or promiscuous bacterial hosts as bait to enrich a diverse range of phages from water samples^10,11^. Additionally, phages must be produced to high titres for clinical use^12,13^. As phage host range narrows, more hosts are required to propagate and biobank a broad range of phages to sufficient titres, and optimisation of host growth parameters suffers combinatorial expansion. Therefore, phage isolation and production efficiency depend on bacterial host promiscuity, which is constrained by anti-phage defence systems^14,15^.

Type I restriction-modification (R-M) systems are the second most common *P. aeruginosa* anti-phage defence system, present in 49% of genomes^16^. R-M systems discriminate between foreign and self-DNA based on methylation status, cleaving phage DNA while protecting the host ^17^. Type I R-M systems consist of three subunits: HsdS, HsdM and HsdR ^18^. One HsdS and two HsdM subunits form a methyltransferase, which methylates the complementary strand at hemi-methylated recognition sites, protecting host DNA from restriction. Adding two HsdR subunits to the methyltransferase forms a restriction endonuclease, which binds to unmethylated recognition sites in foreign DNA and acts as an ATP-dependent motor, translocating DNA and forming a double-stranded break several thousand base-pairs away from the recognition site^19^. Type I R-M systems significantly reduce phage proliferation efficiency^20^, thereby limiting host promiscuity.

Phages can escape restriction by methylating their recognition sites, either by using the host or their own methyltransferase, to produce methylated progeny that are protected in future hosts^21,22^. Additionally, methylation status can affect gene expression, influencing phage virulence, packaging and protein expression ^23,24^. Therefore, while the restriction endonuclease inhibits phage proliferation, the methyltransferase can facilitate the production of methylated phages that are resistant to recognition and cleavage ^21^.

Previously, a *Streptococcus pneumoniae* strain lacking type I R-M systems was used as a permissive host to isolate *Streptococcus* phages^20^. One type I R-M system has been identified in the *P. aeruginosa* laboratory strain PAO1, which is routinely used as a phage enrichment host^25,26^. Here, we sought to construct a more promiscuous host for *Pseudomonas* phage isolation by deleting *hsdR* from PAO1, eliminating the restriction endonuclease while maintaining the methyltransferase. We then evaluated whether using Δ*hsdR* as the bacterial host improved the efficiency of phage propagation and isolation from freshwater and wastewater samples.

## Methods

### Constructing ΔhsdR

An unmarked deletion of *hsdR* (*PA2732*) was generated by two-step allelic exchange with a suicide vector^27^ (strains, plasmids and primers listed in tables 1-3). Approximately 500bp sequences flanking *hsdR* were amplified, joined together by overlap extension PCR, and ligated into the plasmid pEX19gm. Modified plasmids were cloned in *Escherichia coli* DH5α and transformed into *E. coli* Rho3 for conjugation into PAO1 on SOB agar (1.5% bacteriological agar, 20 mM bacto-tryptone, 5 mM bacto-yeast extract, 0.5 mM NaCl, 2.5 mM KCl, 10 mM MgCl_2_). Double recombinants were obtained by counter-selective plating on LB containing 100 mg/mL gentamycin and 5% sucrose. The 1,000 bp region surrounding *hsdR* was PCR amplified and Sanger sequenced to confirm the mutation.

**Table 1.** The strains used in the study.

| Strain | Description | Source |
| --- | --- | --- |
| *Pseudomonas aeruginosa* PAO1 | Wildtype *P. aeruginosa* PAO1 | University of Exeter |
| *Pseudomonas aeruginosa* PAO1 Δ*hsdR* | PAO1 with an unmarked deletion of *hsdR* | This study |
| *Escherichia coli* DH5α | *E. coli* strain used for plasmid cloning | New England Biolabs |
| *Escherichia coli* Rho3 | *E. coli* strain used for plasmid conjugation | Lopez et al (2009) <sup>53</sup> |

**Table 2.** The plasmids used to generate the unmarked deletion of *hsdR*.

| Plasmid | Description | Accession no. | Source |
| --- | --- | --- | --- |
| pEX19gm | Allelic exchange vector encoding gentamycin resistance (*aaC1*) and sucrose sensitivity (*sacB*) | KM887142 | Hoang et al (1998) <sup>54</sup> |
| pEX19gm-Δ*hsdR* | Modified pEX19gm containing ~500 bp regions that flank *hsdR* | | This study |

**Table 3.**
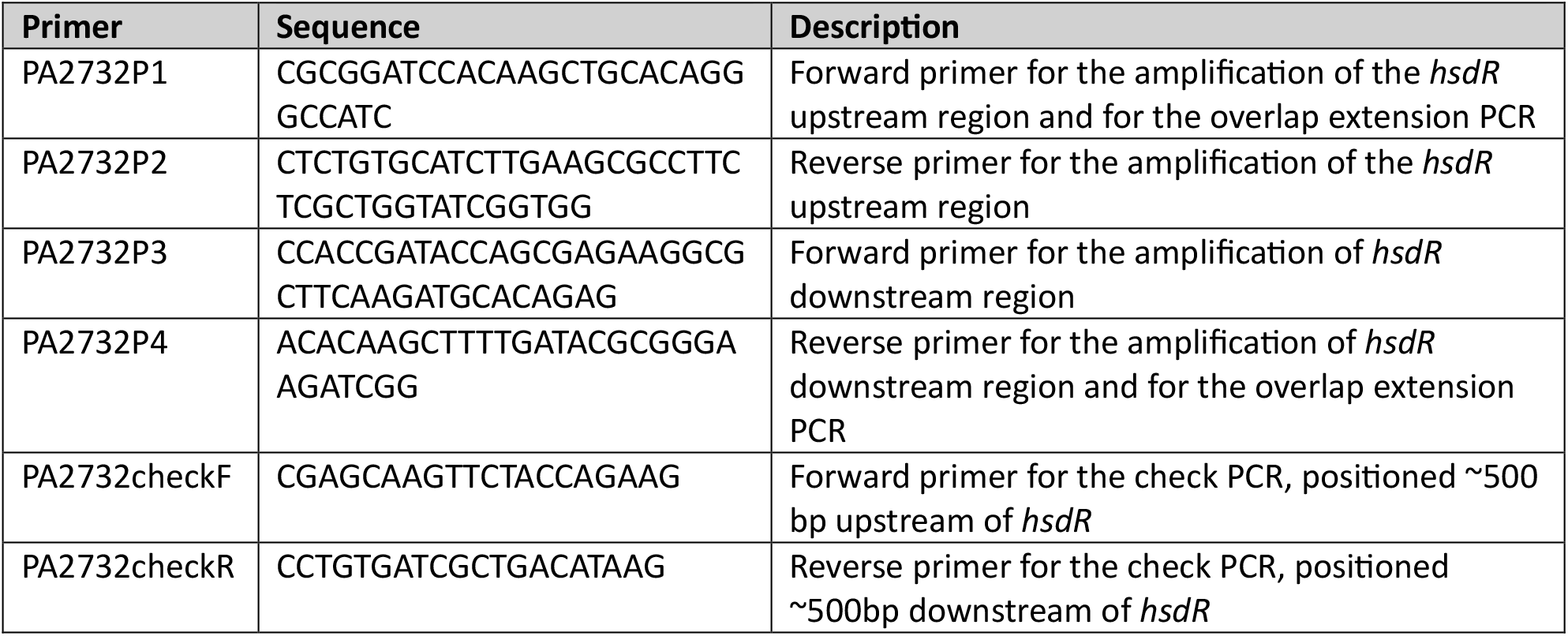
The primers used to generate the unmarked deletion of *hsdR*.

### Evaluation of fitness defects via growth curves

Growth curves were generated to determine whether deleting *hsdR* caused fitness defects in the absence of phage. Cultures of Δ*hsdR* and wildtype PAO1 were grown from single colonies in 10 mL of LB (37 °C, 200rpm) for 3 hours, and then diluted to a starting OD_600_ of 0.1. 100 µL of each culture was added to six replicate wells of a 96-well plate and optical density was recorded every 15 minutes for 16 hours in a plate reader (Tecan Sunrise). The intrinsic growth rate (r), carrying capacity (K) and doubling time (DT) were calculated using Growthcurver (V0.3.1)^28.^

### Evaluation of promiscuity via spot assays

The ability of 86 *P. aeruginosa* phages from the Citizen Phage Library (CPL)^10^ to propagate on Δ*hsdR* and wildtype PAO1 was determined via spot assays^29^. Δ*hsdR* and wildtype PAO1 cultures were grown from single colonies in 10 mL of LB (37 °C, 200rpm) until mid-logarithmic phase (OD_600_ = 0.6). 2 mL of culture was mixed with 6 mL of top agar (0.65% w/v bacteriological agar, 10 mM MgCl_2_, 10 mM CaCl_2_) and poured over a 60 mL bottom agar plate (1% w/v bacteriological agar, 10 mM MgCl_2_, 10 mM CaCl_2_). Phage lysate was serially diluted with LB to a maximum dilution of 10^−6^. A 5 µl spot of each dilution was spotted onto the top agar and incubated overnight at 37 °C. The efficiency of plating (EoP) was calculated for each phage as the proportion of plaque forming units (PFU)/mL between Δ*hsdR* and wildtype PAO1^30^.

### Evaluation of promiscuity via phage isolation

Δ*hsdR* and wildtype PAO1 were used as enrichment hosts for 190 freshwater and 95 wastewater samples. Environmental samples were provided to the CPL by citizen scientists and stored in glass amber vials at 4 °C. Wastewater samples were provided by the Environment Agency and stored in 100 µL aliquots at -20 °C. Following the CPL workflow, reported previously^10^, samples were enriched by two rounds of incubation with the bacterial host, followed by 0.22 µm-filtration. 2 µL of each enriched sample was spotted onto a host bacterial lawn, incubated overnight at 37 °C, and examined for zones of lysis.

### Phage DNA extraction, sequencing and analysis

Agar cores were taken from zones of lysis and transferred to 100 µL of SM buffer (50 mM Tris-HCl (pH 7.5), 0.1 M NaCl, 8 mM MgSO_4_). 50 µL of SM buffer containing phage and 500 µL of overnight bacterial host culture were added to two falcon tubes containing 20 mL of sLB and incubated overnight at 37 °C ^10^. The two 20 mL cultures were combined and centrifuged at 10,000 × g for 30 minutes at 4 °C. 30 mL of supernatant was passed through a 0.22 µm syringe filter and treated with 5 µg /mL DNase I (Roche, Merck, Darmstadt, Germany) and 10 µg /mL RNase A (Invitrogen, ThermoFisher Scientific, Waltham, MA, USA) for 30 minutes at 37 °C. 10% *w/v* polyethylene glycol 8000 (PEG8000) and 1 M NaCl were added, mixed by inversion until dissolved, and incubated overnight at 4 °C. Precipitated phages were pelleted by centrifugation at 10,000 × g for 30 minutes and resuspended in 1 mL SM buffer. DNA was extracted using the Norgen Phage DNA Isolation Kit (Norgen Biotek, Thorold, ON, Canada) and sequenced at the University of Exeter using 2 × 150bp paired end reads on an Illumina NovaSeq SP platform.

Phage genomes were assembled and annotated as follows (full pipelines are available here: https://github.com/citizenphage/protocols/tree/main/SOPs/Assembly-and-annotation): Reads were QCed with FASTP (v. 0.24.1) with the following settings (−-dedup --dup_calc_accuracy 6 --length_required 30 –correction), and high-quality reads were mapped against the genome of the propagation host with minimap2 (v. 2.26) to remove any residual host DNA. Unmapped reads were subsampled to 500-fold coverage with shovill (v. 1.1.0) and assembled with unicycler (v. 0.5.0). Assembly graphs were manually inspected to confirm single circular genomes, with branches of up to 3 bp manually resolved by selection of the most abundant node. Annotation of phage genomes was performed with pharokka (v. 1.7.1) followed by phold (v. 0.1.3).

To quantify the number of HsdR restriction sites in each phage genome, genomic sequences were screened in ipython for non-overlapping recognition sites using the regular expression: “GATC.{6}GTC”.

### Statistical methods

Welch two sample t-tests were used to determine whether the mean growth rate, carrying capacity and doubling time differed significantly between Δ*hsdR* and wildtype PAO1 growth curves. Mean efficiency of plating was bootstrapped 10,000 times with replacement to determine whether it was significantly greater than one. Two-tailed Fisher’s exact tests were used to determine whether the proportion of samples that yield phages differs significantly on Δ*hsdR* and wildtype PAO1.

## Results

### Deleting hsdR posed no fitness cost to PAO1 in the absence of phage

Successful in-frame deletion mutants were identified by a check PCR amplifying the region flanking *hsdR* (supplementary figure 1). Sanger sequencing of the PCR product confirmed the deletion (supplementary material). Growth curves showed that Δ*hsdR* incurred no fitness defects in the absence of phage (supplementary figure 2); there was no significant difference in intrinsic growth rate (r), carrying capacity (K) or doubling time (DT) between Δ*hsdR* and wildtype PAO1 (supplementary table 1).

### ΔhsdR increased the efficiency of phage proliferation

Spot assays revealed that the mean phage titre of 86 *P. aeruginosa* phages was 2.15-fold higher on Δ*hsdR* than wildtype PAO1, with a mean efficiency of plating (EoP) of 4.31 (figure 1A). When bootstrapped 10,000 times with replacement, the mean EoP was always greater than one, indicating that phage proliferation was significantly more efficient on Δ*hsdR* than wildtype PAO1 (figure 1B, *p*<0.0001). Two phages, CPL00163 and CPL00284, did not form plaques on wildtype PAO1 but did on Δ*hsdR,* with a titre of 6.40 × 10^7^ and 8.20 × 10_5_ PFU/mL respectively, indicating that HsdR completely inhibits their propagation. The number of HsdR recognition sites (GATC(N) _6_GTC)^26^ found within CPL00163 (7) and CPL00284 (6) sat within the range of the number of sites found in 16 other phages that could infect both Δ*hsdR* and wildtype PAO1 (3-7, mean=4.6±0.62 95% CI), suggesting that the prevalence of recognition sites does not determine phage specificity between Δ*hsdR* and wildtype PAO1.

**Figure 1:**
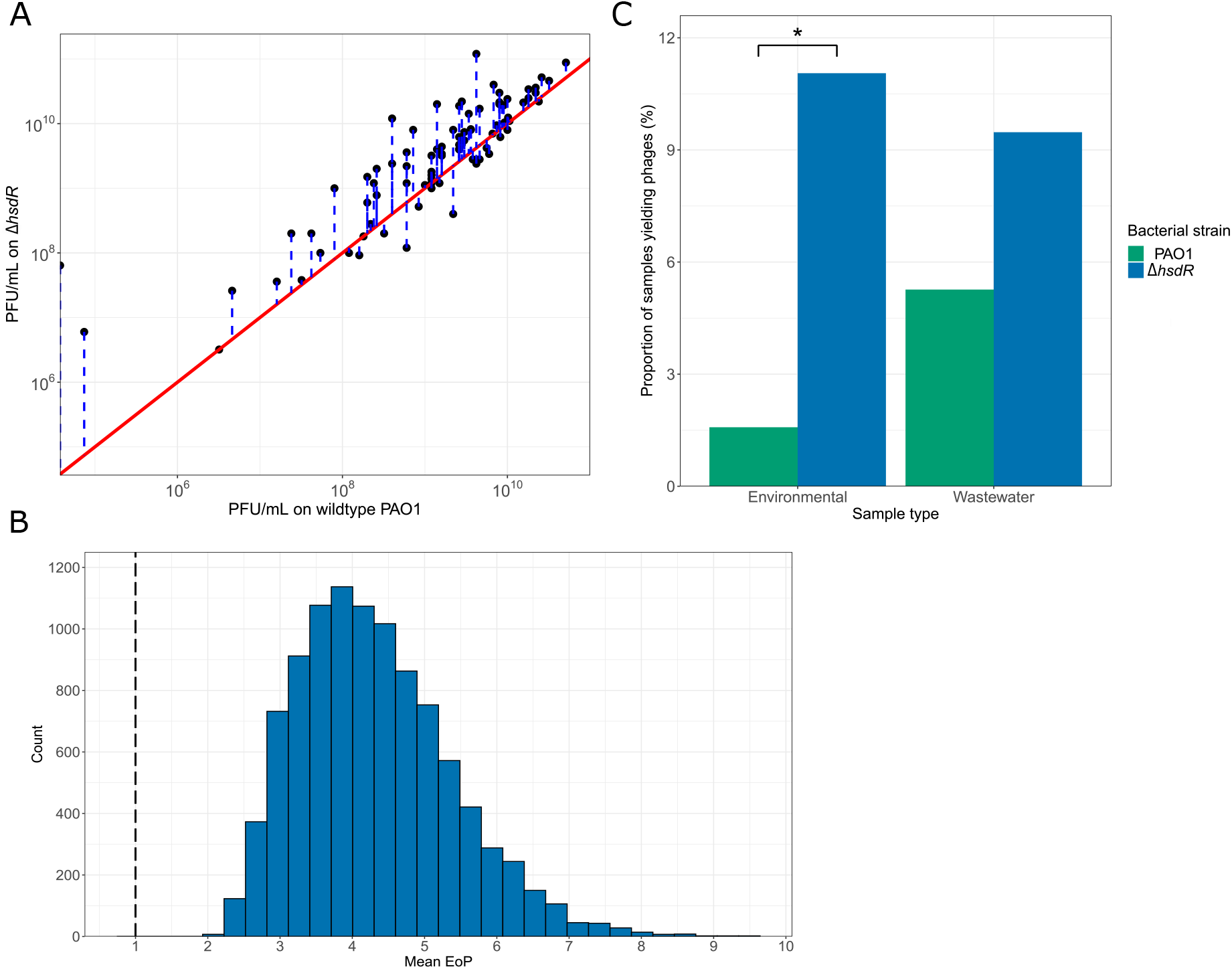
**[A]** Comparison of Concentration (PFU/mL) of each phage on wildtype PAO1 against its concentration on the Δ*hsdR* mutant. The distance (blue) between each data point and the identity y=x (red) represents the difference in PFU between the strains. **[B]** Bootstrapping of the mean efficiency of plating (EoP) of phages on Δ*hsdR* compared to wildtype PAO1 10,000 times with replacement. All bootstrapped means are greater than one, indicating that PFU is consistently higher across phages on PAO1 Δ*hsdR* than the wildtype PAO1. **[C]** The proportion of environmental (green) and wastewater (blue) samples that yielded phages when enriched with wildtype PAO1 and Δ*hsdR* as hosts. Two-tailed Fisher’s exact tests were used to determine whether there was a significant difference in the proportion of water samples yielding phages between bacterial host strains (\**p*<0.001).

### ΔhsdR increased the phage isolation yield from freshwater samples

In total, 24 and 14 phages were isolated from 190 freshwater and 95 wastewater samples, respectively. There was a significant 7.00-fold increase in the proportion of freshwater samples that yielded phage on Δ*hsdR* compared to wildtype PAO1 (Fisher’s exact test, *p<*0.001, OR = 0.130, 95% CI = [0.0243, 0.446]) (figure 1C). Curiously, this was not observed with wastewater samples (Fisher’s exact test, *p=*0.406, OR = 0.533, 95% CI = [0.135, 1.854]) (supplementary tables 2 & 3). Only 1 of the 13 wastewater samples and 1 of the 23 freshwater samples that yielded phages did so on both Δ*hsdR* and wildtype PAO1, indicating that *Pseudomonas* phage abundance and diversity within and between sample types were highly variable.

## Discussion

Effectively providing phages for clinical use requires comprehensive libraries for rapidly identifying specific phages and streamlined methods for producing sufficiently high titres^9,10,12^. The efficiency of constructing diverse libraries and propagating a broad range of phages depends on the promiscuity of the bacterial host. Here, we created a more promiscuous *P. aeruginosa* host by deleting the defensive restriction endonuclease HsdR.

Phage propagation was 2.15-fold more efficient on Δ*hsdR* than wildtype PAO1, aligning with previous findings that restriction-modification systems significantly limit phage proliferation^20,21^. One study reported a 10,000-fold decrease in phage SpSL1 proliferation when the phase-variable type I R-M system SpnIV was expressed in *S. pneumoniae* ^20^. Here, we found that deleting the HsdR subunit alone was sufficient to increase phage proliferation. This deletion eliminates phage DNA cleavage by the restriction endonuclease but maintains the methyltransferase, such that phage progeny may still be methylated and protected in future hosts^21,22^. Similarly, a previous study found that increasing the ratio of methyltransferases to restriction endonucleases in an *E. coli* type II R-M system abolished protection against phage^21.^

Two phages (CPL00163 and CPL00284) could infect Δ*hsdR* but not wildtype PAO1. Thus, Δ*hsdR* permits infection by some phages that are otherwise inhibited by restriction. CPL00163 and CPL00284 possessed a similar number of recognition sites to other phages that can infect wildtype PAO1, indicating that a high frequency of recognition sites is not the reason that these phages are particularly sensitive to restriction ^31.^ Alternatively, they might lack anti-restriction strategies that other phages possess ^32,33^. For instance, some phages avoid recognition by acquiring methylated or non-canonical bases in their recognition sites ^24,34,35^, or by encoding anti-restriction proteins like Ocr, which blocks the endonuclease binding groove ^36^.

Using Δ*hsdR* as the enrichment host increased freshwater enrichment yields sevenfold but had no significant impact on wastewater enrichment yield. Phage concentrations are typically 10-1000 times higher in wastewater than freshwater^37–39^, making wastewater phage enrichments generally more successful^40^. Since Δ*hsdR* enables more efficient phage proliferation than wildtype PAO1, using Δ*hsdR* as the host may increase enrichment yields by improving the chance of recovering low-abundance phages. As a result, eliminating *hsdR* may have a more significant impact on freshwater isolation yields, where phages are typically present at lower concentrations.

The overall success rate of phage enrichment was relatively low compared to previous studies ^11,41,42^. One study achieved a 79.4% success rate for isolating *P. aeruginosa* phages from 20-30 mL samples of unprocessed wastewater^41^. We may have recovered fewer phages from wastewater as our samples were stored at – 20 °C and freeze-thawed, potentially damaging phages^43,44^. Additionally, enriching smaller 1 mL samples may have reduced the success rate per sample, but allowed overall higher throughput. The relatively low chance of yielding a phage from each sample with this high-throughput method may explain why there was little overlap in the samples that yielded phage on Δ*hsdR* and wildtype PAO1, as it was unlikely for the same phage to be isolated more than once.

Since phage proliferation was more than twice as efficient on Δ*hsdR*, more phages could be propagated to sufficient titres for clinical use on this strain. This would reduce the number of specific host strains that need to be tested and optimised, streamlining phage production. To further optimise Δ*hsdR* for propagating phages, secreted virulence factors, such as exotoxin A, could be deleted to expediate downstream purification^45,46^. As freshwater isolation yields were seven times greater on Δ*hsdR*, a broader range of phages could be efficiently obtained for phage libraries using this strain. The recent discovery of phage defence islands in bacterial genomes has uncovered dozens of novel anti-phage defence mechanisms^15,32,47^, which could be future targets for deletion to further increase Δ*hsdR* promiscuity. Beyond defence systems, host promiscuity is constrained by phage receptor specificity^48,49^. Similar deletions of anti-phage defence systems and toxins could be made in other *P. aeruginosa* strains that possess different variations of common phage receptors, such as the LPS and type IV pili^50–52^. Ultimately, this would create a panel of characterised, promiscuous and safe hosts for the streamlined isolation and propagation of a diverse range of phages.

## Supporting information

Phages used in this study

## Data availability

Sanger sequence data for the Δ*hsdR* deletion mutant is provided at https://github.com/citizenphage/Tong-hsdR. Source data and information on the phages used in this study are provided in the supplementary materials. NCBI Accession numbers for phages used in this study are provided in Supplementary Table 4.

## Code availability

The data analysis in this study was performed with R and python code, available here (https://github.com/citizenphage/Tong-hsdR).

## Acknowledgments

The authors would like to acknowledge the contributions of Matt Scurlock and Christian Fitch for their support and training in gene deletions and phage propagation, respectively. The authors would also like to acknowledge Dr. Luis M. Bolaños for his support in data submission.

## Author contributions

E.T. constructed the Δ*hsdR* deletion mutant, performed the data analysis and wrote the manuscript. E.T., K.B. and A.C. performed the spot assays and phage enrichments. B.T. conceived the experimental design and performed the phage genome assemblies. B.T. and S.P. edited the manuscript.

