## Supplementary figure 1 for "Eliminating the type I restriction endonuclease from *Pseudomonas aeruginosa* PAO1 for optimised phage isolation"

**Supplementary materials**

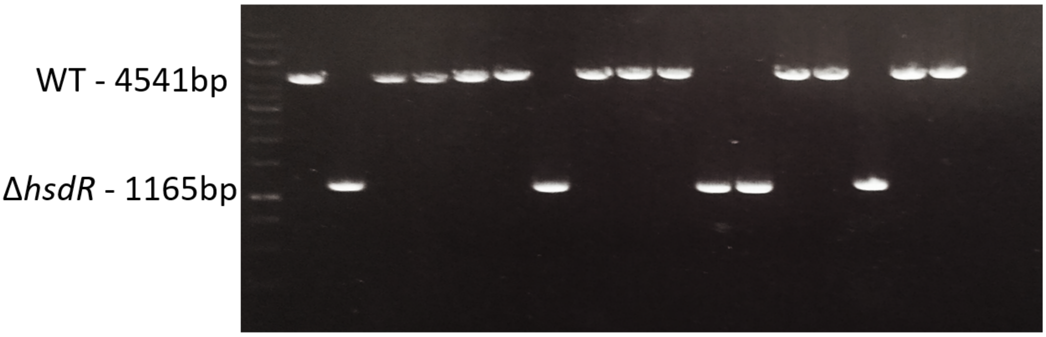


**Supplementary figure 1:** An image of the check PCR products run on a 1% agarose gel. The sequence flanking *hsdR* was PCR amplified to distinguish successful in-frame deletion mutants (1165 bp) from wildtype (WT) revertants (4541 bp).

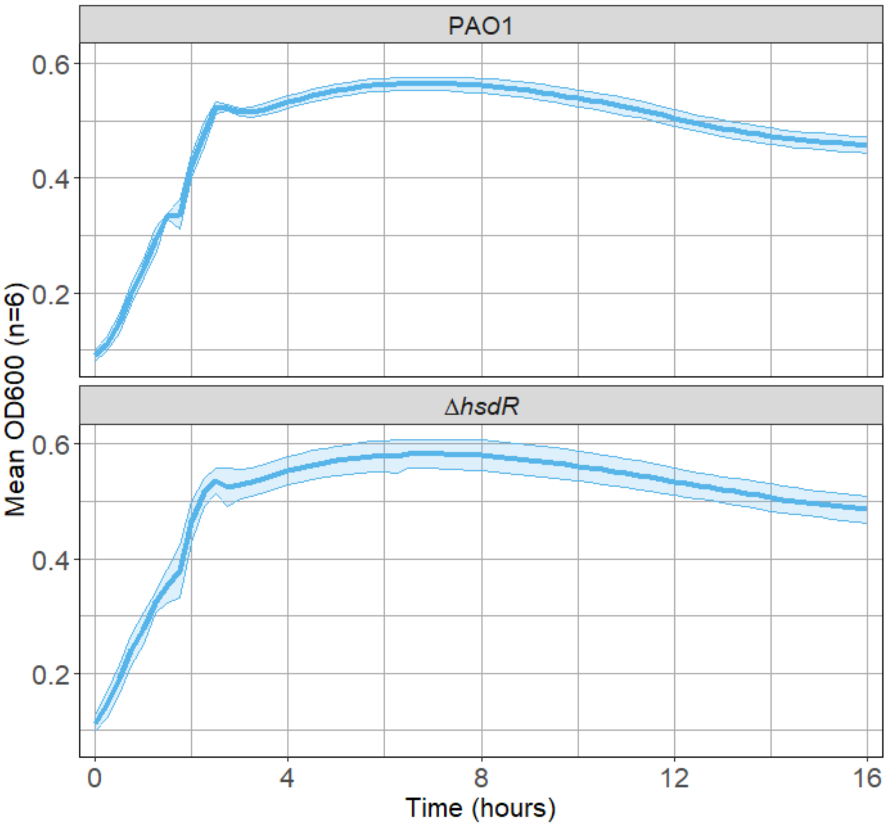


**Supplementary figure 2:** The mean $OD_{600}$ of Δ*hsdR* and wildtype PAO1 over 16 hours at 37 °C. Ribbons represent 95% confidence intervals around the mean.

| | **Carrying capacity (K)** | **Intrinsic growth rate (r)** | **Doubling time (DT)** |
|---|---|---|---|
| **Mean value for PAO1** | 0.394 | 2.51 | 0.277 |
| **Mean value for Δ*hsdR*** | 0.396 | 2.42 | 0.287 |
| **Degrees of freedom (df)** | 6.77 | 6.48 | 6.17 |
| ***p*-value** | 0.822 | 0.242 | 0.233 |

**Supplementary table 1:** The mean carrying capacity (K), intrinsic growth rate (r) and doubling time (DT) of PAO1 and Δ*hsdR* determined using Growthcurver[28]. Welch two sample t-tests were used to determine if the mean growth parameters differed significantly between PAO1 and Δ*hsdR*.

| | **Zone of lysis** | **No zone of lysis** |
|---|---|---|
| **PAO1** | 3 | 187 |
| **Δ*hsdR*** | 21 | 169 |

**Supplementary table 2:** Contingency table for the proportions of freshwater samples that yield phage on PAO1 and Δ*hsdR*

| | **Zone of lysis** | **No zone of lysis** |
|---|---|---|
| **PAO1** | 5 | 90 |
| **Δ*hsdR*** | 9 | 86 |

**Supplementary table 3:** Contingency table for the proportions of wastewater samples that yield phage on PAO1 and Δ*hsdR*
